# Biofilm selection constitutively activates c-di-GMP synthesis by the bifunctional enzyme MbaA

**DOI:** 10.1101/2025.02.13.638143

**Authors:** Øyvind M. Lorentzen, Sören Abel, Pål J. Johnsen, Christopher Frøhlich

## Abstract

Biofilms, in which microbes are encased within a self-produced matrix, represent the primary mode of microbial life. Yet, our understanding of the molecular mechanisms governing biofilm formation remains incomplete, limiting the development of targeted interventions against biofilm-associated infections. By selecting for *Vibrio cholerae* biofilms, we identified mutations in the polyamine-regulated, bifunctional enzyme MbaA that alter its enzymatic activity in metabolizing the conserved second messenger cyclic diguanylate (c-di-GMP). These mutations occur in both the c-di-GMP-producing GGDEF domain and the c-di-GMP- hydrolyzing EAL domain of MbaA. Mutagenesis and enzyme kinetic analyses revealed regulatory crosstalk between these domains, where mutations in either domain enhanced production of c-di-GMP by MbaA while concurrently reducing its hydrolytic activity. In the wild-type enzyme, c-di-GMP metabolism is regulated by specific polyamines. Biofilm-evolved mutations not only shift MbaA’s enzymatic activity but also disrupt this post-translational regulation, rendering the enzyme insensitive to external polyamine signaling. Our findings illustrate how evolutionary pressure can rewire existing enzymes, driving a transition from a planktonic to a biofilm-associated lifestyle.

## INTRODUCTION

The formation of biofilms, microbial communities enclosed in a self-produced matrix, is an essential lifestyle that increases survival and adaptation^1–4^. The protective nature of biofilms complicates the treatment of biofilm-associated infections, often leading to recurrence and therapeutic failure^1,3,5^. Additionally, biofilms play an important role in bacterial adaptation as this lifestyle allows for evolutionary trajectories that differ from those in planktonically-growing bacteria^6–16^. Despite the importance of biofilms, our understanding of the molecular mechanisms that regulate lifestyle transitions from planktonic to biofilm-associated remains incomplete. This limits not only our evolutionary understanding of bacterial behavior but also complicates the design of targeted interventions against biofilm-producing pathogens.

One of the key regulators of biofilm formation in bacteria is the second messenger cyclic diguanylate (c-di-GMP)^17–19^. Typically, low intracellular levels of c-di-GMP are associated with a motile single cell lifestyle, while high levels are associated with biofilm formation^17–19^. The concentration of c-di-GMP is controlled by enzymes that produce or degrade c-di-GMP through three enzyme domains, which are named after conserved sequence motifs within their active sites. Diguanylate cyclases (DGCs), hallmarked by GGDEF domains, catalyze the synthesis of c-di-GMP from GTP. In contrast, phosphodiesterases (PDEs), which contain EAL or HD-GYP domains, catalyze the hydrolysis of c-di-GMP into pGpG or GMP, respectively^17–20^. These domains can also be found together in the same enzyme giving rise to potentially bifunctional c-di-GMP-metabolizing enzymes^17–21^. Due to their importance for bacterial lifestyle regulation, c-di-GMP-metabolizing enzymes are typically under stringent post-translational regulation by either regulatory proteins or sensory domains^17–19,21^. One example of such regulators are polyamines, small molecules such as norspermidine and spermidine that enhance or suppress biofilm formation in multiple bacterial species^22–26^. Disrupting this regulation can alter the cellular equilibrium of c-di-GMP, driving the transition between a motile and a biofilm-associated lifestyle and *vice versa*^11,17–20,27^.

Among the bifunctional c-di-GMP-metabolizing enzymes is MbaA, a membrane-bound enzyme comprising a signaling domain (HAMP) followed by GGDEF and EAL domains^22,28^. Previous studies have demonstrated that loss of *mbaA* enhances biofilm formation in *V. cholerae* and that its activity is under post-translational regulation by the polyamine sensor NspS^22,23,25,28,29^. In its native state, MbaA exhibits PDE activity and hydrolyzes c-di-GMP, thereby inhibiting biofilm formation. Upon norspermidine binding to the polyamine-sensing protein NspS, the NspS-norspermidine complex is believed to interact with MbaA resulting in a shift of its enzymatic activity from degradation to synthesis of c-di-GMP, thereby increasing biofilm formation^22,23,25,30^. In contrast, binding of spermidine to NspS amplifies degradation of c-di-GMP, thereby inhibiting biofilm formation^22,23,25,30,31^.

In this study, we employed experimental evolution of *V. cholerae* biofilms to examine the genetic changes that occur in the c-di-GMP signaling system. We identified mutations in the bifunctional c-di-GMP-metabolizing enzyme MbaA that induced transition to a biofilm- associated lifestyle. By combining biofilm measurements, mutagenesis studies and *in vitro* enzyme kinetics, we uncover that biofilm-evolved mutations in MbaA reduce the PDE activity of the EAL domain while simultaneously increasing c-di-GMP production by constitutively activating the GGDEF domain. In addition, these mutations decouple MbaA from post- translational regulation by NspS, thereby desensitizing it to external signals. Ultimately, this study provides mechanistic insight into how bacteria can leverage the evolutionary potential of bifunctional c-di-GMP-metabolizing enzymes to facilitate adaptation.

## RESULTS

### Selection for biofilm formation results in mutations in *mbaA*

To select for increased biofilm formation, we propagated biofilm pellicles at the air-liquid interface of static *V. cholerae* C6706 cultures every 24 hours for 8 days (Fig. 1a). We evolved six biofilm populations and, as a control, two planktonic populations sampled from the culture underneath the pellicles and investigated the ability of these populations to produce biofilms by crystal violet staining. The planktonically-evolved populations exhibited only a marginal increase in biofilm formation (2-3-fold), which was not statistically significant, compared to the parental strain (Fig. 1b, Tab. S3, one-way ANOVA, *P* >0.05). In contrast, the biofilm-evolved populations displayed a significant 8 to 11-fold increase in attached biomass (Fig. 1b, Tab. S3, one-way ANOVA, *P* < 0.01). In addition, the biofilm-evolved populations produced pellicles with a wrinkly morphology (Fig. 1c), a trait associated with hyper-biofilm phenotypes ^32–35^. In contrast, the parental strain and planktonically-evolved populations produced smooth pellicles. To identify the underlying genetic changes driving the observed increase in biofilm formation, we subjected all evolved populations to whole genome sequencing (File S1). The planktonically-evolved populations primarily accumulated mutations in genes related to flagellum biosynthesis and quorum sensing (*flrAB*, *motX*, and *luxO*), which have previously been described to occur during propagation in liquid culture. (Fig. 1d, File S1)^36–38^. In addition, the planktonic-evolved populations acquired mutations in the GGDEF domain of a DGC (VCA0697), although at a very low frequency (<2%). In contrast, all biofilm-evolved populations exhibited strong signs of convergent evolution, with high frequencies of mutations occurring in *mbaA* (33.7% to 82.8%), which encodes a bifunctional c-di-GMP metabolizing enzyme (Fig. 1d-e). In addition, a subset of the biofilm-evolved populations acquired mutations in the global regulator *cytR* and in the transport system *ugpE* (Fig. 1d).

**Figure 1:**
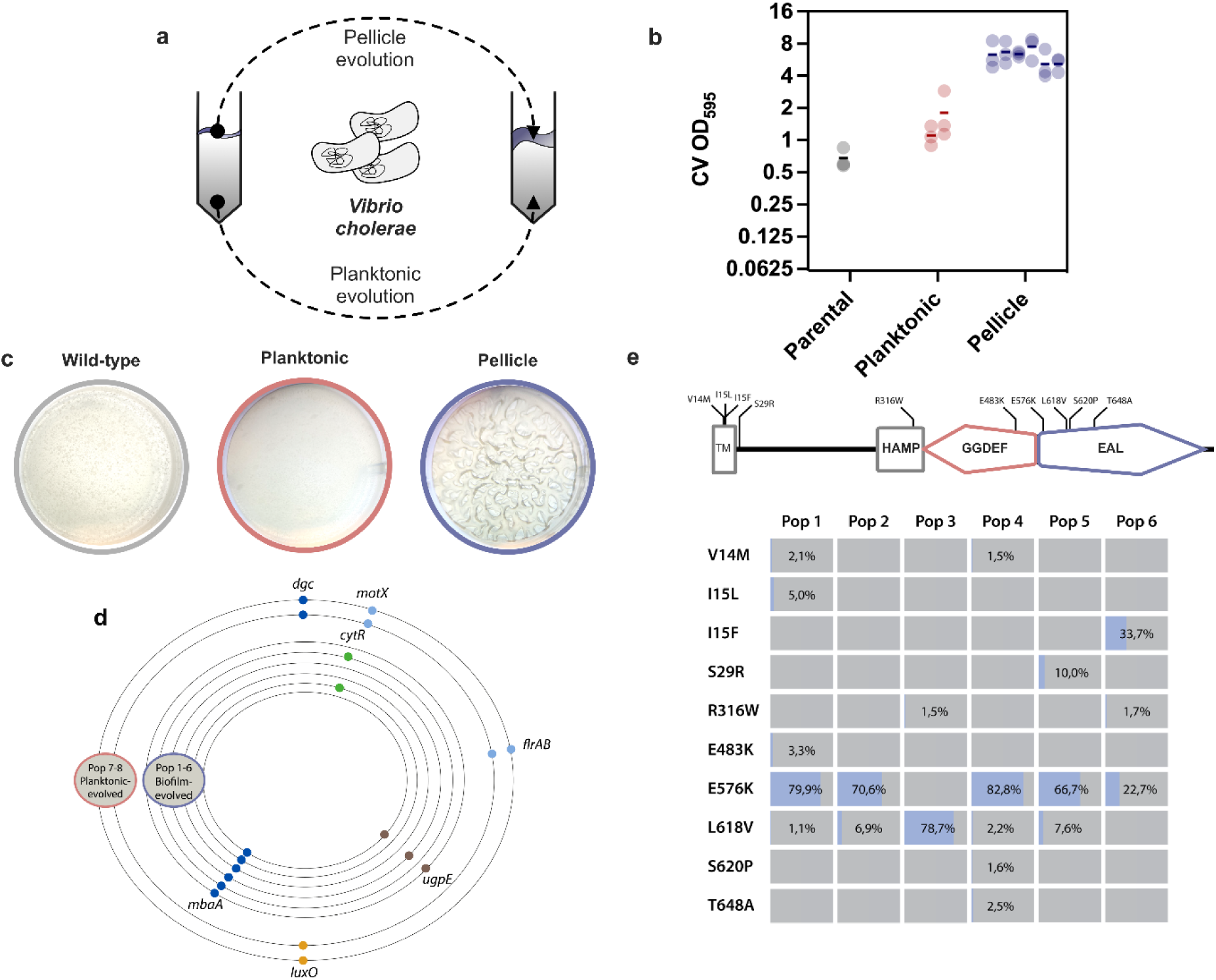
Evolution of *V. cholerae* biofilm pellicles selected for increased biofilm formation and mutations in MbaA. **a.** Schematic representation of experimental evolution protocol. The biofilm pellicle from six statically grown *V. cholerae* cultures and planktonic bacteria from the liquid phase from two statically grown *V.* cholerae cultures were propagated every 24 hours for 8 days. **b**. Biofilm formation of parental *V. cholerae*, planktonically- (*n* = 2) and biofilm-evolved populations (*n* = 6). Biofilm production was quantified by crystal violet (CV) biofilm formation assays, measuring absorbance at 595 nm of retained CV. Dots represent one biological replicate and the error bars represent the standard error of the mean. Statistical significance was tested with one way ANOVA followed by Dunnett test to correct for multiple testing. See Tab. S3 for *P*-values of all evolved populations. Each population was tested with three biological replicates. **c.** Representative images of biofilm pellicles morphologies from the parental strain, planktonically- or biofilm-evolved populations after 24 h of static growth. **d.** Mutations found after biofilm selection mapped to the chromosome. Each ring represents the combined genomes of one evolved population. Biofilm-evolved populations 1 to 6 (blue), planktonic-evolved populations 7 to 8 (red). Dots indicate mutations discovered in at least two populations (File S1 contains complete list of all identified mutations). Dots are colored depending on the function/cellular process: dark blue = c-di-GMP signaling, brown = transportation, green = transcriptional regulation, light blue = flagellum biosynthesis/motility and orange = quorum sensing. **e.** Frequency, position, and amino acid changes of all identified mutations in MbaA within each of the biofilm-evolved populations.

In total, we identified 10 different mutations in *mbaA* distributed over the N-terminal transmembrane domain (V14M, I15L, I15F, S29R), the regulatory HAMP-domain (R316W), the GGDEF domain (E483K) and EAL domain (E576K, L618V, S620P, T648A) (Fig. 1e). The most frequent mutation was E576K, located in the EAL domain, which was present in five out of six biofilm-evolved populations at frequencies between 22.7 to 82.8%. Aligning MbaA against a selection of GGDEF-EAL domain containing proteins indicated that several of the mutations in the GGDEF and EAL domain (E483K, E576K, L618V, S620P and T648A) were in close proximity to putative conserved amino acid residues (Fig. S1-2), thereby potentially affecting the enzymatic function of MbaA^18,20,39–43^.

### Interplay between the GGDEF and EAL domain induces a hyper-biofilm phenotype

Whole genome sequencing identified mutations in *mbaA* in all of the biofilm-evolved populations, indicating that *mbaA* was under strong selection during biofilm evolution (Fig. 1d, File S1). Since evolved populations instead of isolated clones were sequenced, it could not be excluded that multiple mutations within *mbaA* were required to increase biofilm formation. Therefore, two clones were isolated from the evolved populations containing either a single E576K or a single L618V mutation in MbaA. These represent the most prevalent mutations in the EAL domain. Additionally, to cover the mutation identified in the GGDEF domain, we engineered E483K in the parental strain background (Fig. 1e, Tab. 1). Assessing biofilm formation, strains containing the isolated mutations in the GGDEF or EAL domain had significantly (one-way ANOVA, *P* <0.0001) increased biofilm formation by 12- to 15-fold compared to the parental strain (Fig. 2a, Tab. 1, Fig. S3). Like the biofilm-evolved population, these individual mutants also produced wrinkly biofilm pellicles (Fig. 2b). Our results show that a single amino acid change in *mbaA* is sufficient to induce a hyper-biofilm phenotype, exceeding the observed biofilm forming capacity of the evolved populations containing a mix of mutants.

**Figure 2:**
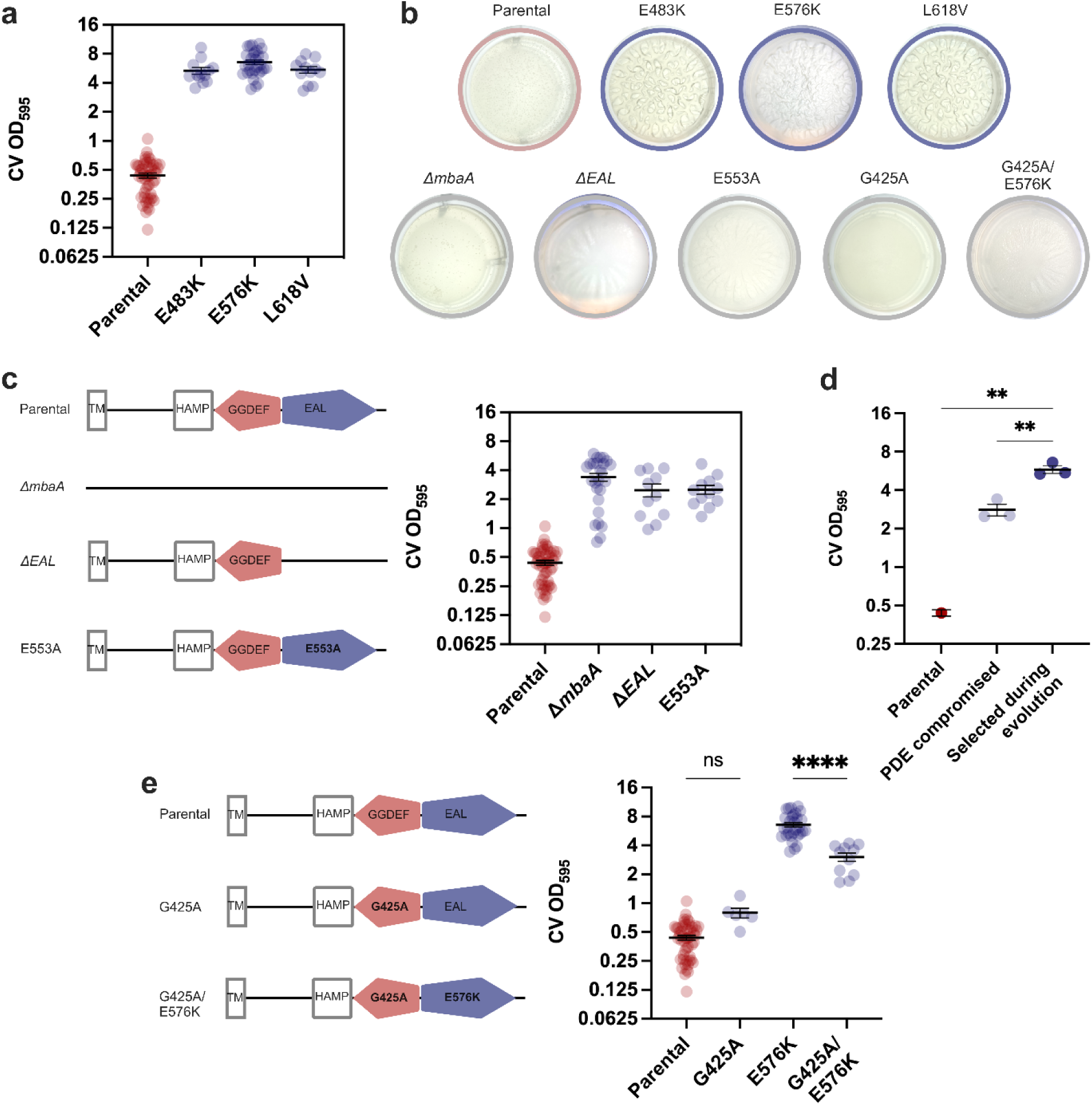
Hyper-biofilm phenotype depends on the crosstalk between GGDEF and EAL domain. **a.** Biofilm formation of parental *V. cholerae* and individual MbaA mutants (E483K, E576K and L618V). All MbaA mutants displayed significantly increased biofilm forming capacity (*P* < 0.0001, one-way ANOVA followed by Dunnett’s T3 multiple comparisons tests, Tab. 1) compared to the parental strain. **b.** Only the hyper-biofilm variants E483K, E576K and L618V produce wrinkly pellicles. Representative images of the morphology of biofilm pellicles formed by *V. cholerae* producing either wild-type or MbaA mutants. **c.** MbaA mutants with compromised PDE activity (Δ*mbaA*, Δ*EAL* and E553A) exhibited significantly increased biofilm forming capacity (*P* < 0.01, one-way ANOVA followed by Dunnett’s T3 multiple comparisons tests, Tab. 1) compared to the parental strain. **d.** MbaA mutants selected for by biofilm evolution (E483K, E576K, L618V) demonstrated significantly increased biofilm formation, compared to mutants with compromised PDE activity (Δ*mbaA*, Δ*EAL* and E553A; **; one-way ANOVA, *P* < 0.01, see Tab. S4 for *P*-values of all individual mutants). **e.** While G425A did not alter biofilm formation compared to wild-type MbaA (ns; one-way ANOVA, *P* > 0.05), introducing G425A into E576K led to a significant decrease in biofilm formation compared to E576K (****; one-way ANOVA, *P* < 0.0001, see Tab. S4 for *P*-values of all individual mutants). For all panels, biofilm production was quantified by crystal violet (CV) biofilm assays. Each dot represents one biological replicate. Error bars represent standard error of the mean.

**Table 1.** Biofilm formation of *V. cholerae* producing different MbaA variants.

| Variants | Biofilm formation<br>OD<sub>595</sub><sup>a, e</sup> | Relative biofilm<br>formation<sup>b</sup> | *P* values<sup>c</sup> | *N*<sup>d</sup> | Pellicle<br>morphology |
| --- | --- | --- | --- | --- | --- |
| Parental | 0.44 ± 0.03 | 1 | - | 51 | Smooth |
| E483K | 5.35 ± 0.43 | 12.2 | <0.0001 | 12 | Wrinkly |
| E576K | 6.58 ± 0.36 | 15.1 | <0.0001 | 30 | Wrinkly |
| L618V | 5.46 ± 0.42 | 12.5 | <0.0001 | 12 | Wrinkly |
| ΔmbaA | 3.39 ± 0.32 | 7.8 | <0.0001 | 26 | Smooth |
| ΔEAL | 2.49 ± 0.38 | 5.7 | 0.0085 | 11 | Smooth |
| E553A | 2.52 ± 0.27 | 5.8 | 0.0003 | 12 | Smooth |
| G425A | 0.80 ± 0.09 | 1.8 | 0.1459 | 6 | Smooth |
| G425A/E576K | 3.00 ± 0.30 | 6.9 | 0.0002 | 11 | Smooth |

<sup>a</sup> Absorbance measured at 595 nm for dissolved biofilms stained with crystal violet (CV) in 96-well plates. Errors reported represent the standard error of the mean.
<sup>b</sup> Relative biofilm formation normalized to parental *V. cholerae* C6706.
<sup>c</sup> Brown-Forsythe one-way ANOVA to test for differences in biofilm formation compared to parental *V. cholerae* ( $\alpha=0.05$ ) and followed by Dunnett T3 test to correct for multiple testing; F (DFn, DFd) = 69.11 (8.000, 98.91); overall *P* value < 0.0001.
<sup>d</sup> Sample size (biological replicates) tested to determine the attached and crystal violet stained biomass at OD<sub>595</sub>.
<sup>e</sup> Results are visualized in Fig. S3.

Since MbaA primarily acts as a PDE in its native state^22,23,25,30^, we aimed to investigate the role of the EAL domain in regulating biofilm formation. For that, a whole-gene knockout mutant (Δ*mbaA*), a MbaA mutant lacking the EAL domain (Δ*EAL*) and an active site mutant with impaired PDE activity (E553A, creates an AVL-motif in MbaA) was constructed)^22,25,39–43^.

*V. cholerae* harboring either of these three genetic changes exhibited a statistically significant (one-way ANOVA, *P* <0.001) increase in biofilm production, which was 6 to 8-fold higher than the parental strain (Fig. 2c, Tab. 1). However, biofilm formation by these mutants was significantly lower (one-way ANOVA, *P* <0.01) relative to the individual E483K, E576K, and L618V variants (Fig. 2d, Tab. 1, Tab. S4, Fig. S3). In addition, none of these mutants produced wrinkly biofilm pellicles (Fig. 2b). While the removal of either *mbaA* or the EAL domain as well as compromising the active site of the EAL domain (E553A) increased biofilm formation, it was insufficient to reproduce the hyper-biofilm phenotype that the evolved variants exhibited (E483K, E576K, L618V).

Next, we investigated the dependency of the hyper-biofilm phenotype on the GGDEF domain. In the parental strain, we engineered a mutation in the conserved GGDEF motive of MbaA, known to reduce DGC activity in other DGCs, by replacing glycine in position 425 with an alanine (G425A)^35^. Compromising the conserved GGDEF motive (G425A) resulted in similar levels of biofilm compared to the parental strain (Fig. 2e, Tab. 1). This was expected, since MbaA functions as a PDE under the tested growth conditions, impairing DGC activity should not affect biofilm formation significantly (one-way ANOVA, *P* = 0.15). Next, we combined G425A with the E576K mutation. The combined mutations (G425A/E576K) led to a significant 2.2-fold decrease (one-way ANOVA, *P* <0.0001, Tab. S4) in biofilm formation compared to E576K and loss of the wrinkly pellicle phenotype (Fig. 2b, Fig. 2e, Tab. 1, Fig. S3). Thus, the hyper-biofilm phenotype induced by the E576K mutation in the EAL domain depends on the GGDEF domain and its ability to produce c-di-GMP. Altogether, our mutagenesis study suggests that the hyper-biofilm phenotype is the result of a changed interactions between the GGDEF and EAL domain that activates the DGC activity of MbaA.

### Mutations increase the DGC activity and impair the PDE activity of MbaA

Our mutagenesis study indicated that the observed hyper-biofilm phenotype depends on the interplay between the GGDEF and EAL domain (Fig. 2, Tab. 1). To visualize the location of these mutations within the structure of MbaA, a homology model was created (Fig. 3a). The E576K and L618V mutations within the EAL domain were located in the active site and at the interface between the GGDEF and EAL domain (Fig. 3a-b). Our model suggests that E576 could be involved in the binding of c-di-GMP and likely interacts with R437 within the GGDEF domain through electrostatic interactions (Fig. 3b). The structural role of L618 remains elusive, but its proximity to multiple conserved residues in the active site of the EAL domain suggests a potential effect on the activity of MbaA (Fig. S1). The E483K mutation in the α4-helix of the GGDEF domain is outside of the active site. Interestingly, this region has been described to be involved in electrostatic interdomain interactions in other GGDEF-EAL domain proteins^44,45^. Finding mutations that are located within the interdomain interface is in line with our hypothesis that they may affect crosstalk between domains and alter the enzymatic activity of both enzyme domains.

**Figure 3:**
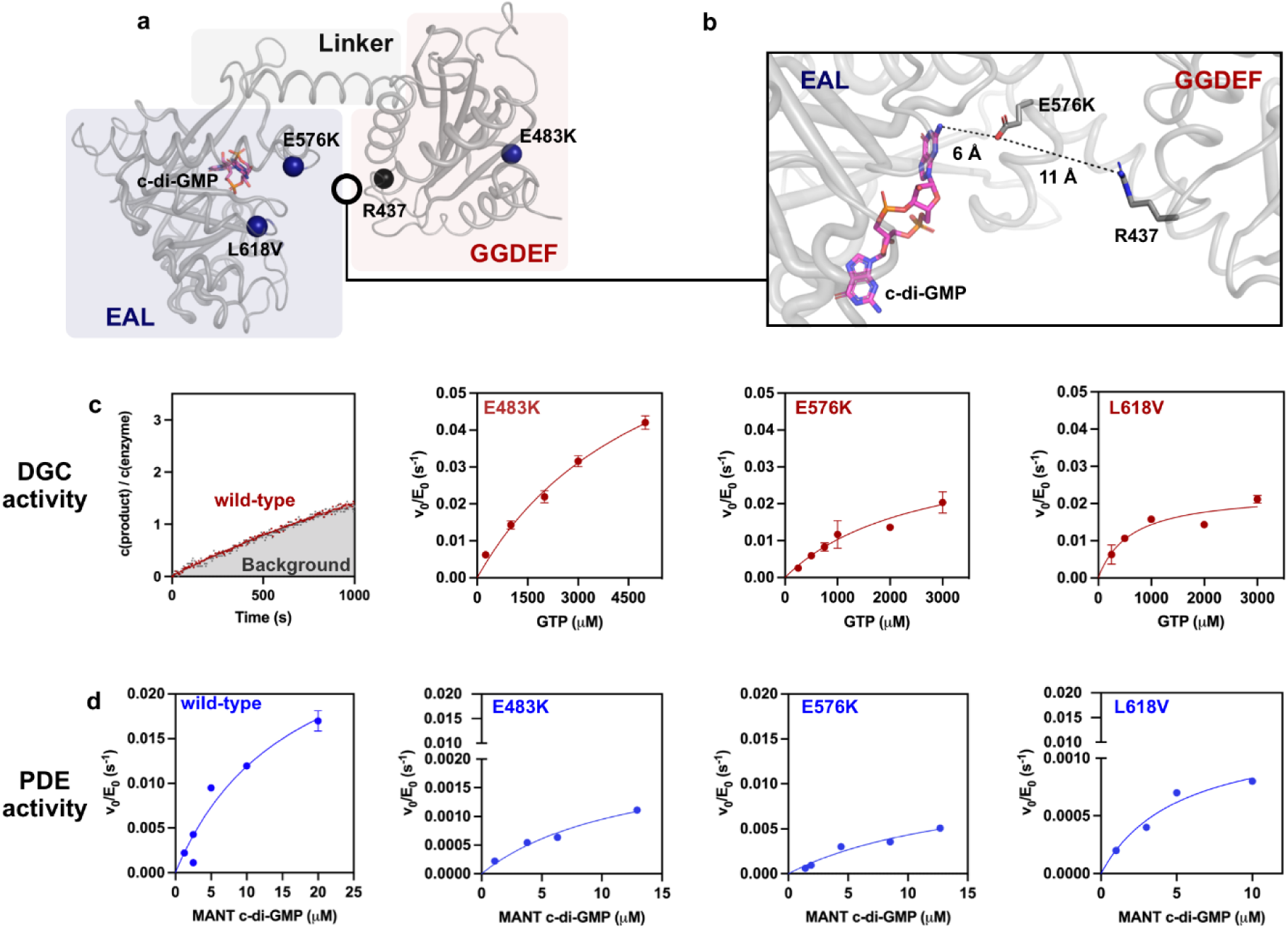
Structural model and enzyme kinetics of MbaA and evolved mutants. **a.** Modelled structure of MbaA with c-di-GMP-bound EAL domain (blue), linker (grey) and GGDEF domain (red). Investigated mutations within the domains are shown as blue spheres. R437, the potential interaction partner of E576 the GGDEF domain, is shown with a black sphere. **b.** Close-up view of the molecular environment surrounding the most frequently selected mutation, E576K. The mutated residue and its potential interaction partner (R437) are shown in the homology model. Potential electrostatic interactions are indicated by dotted lines and the modelled distance between residues is displayed. **c.** DGC enzymatic kinetics by determination of released Pi from the reaction of GTP to c-di-GMP. For the wild-type enzyme (red line), no pyrophosphate release was detectable above the background reaction (grey dots) when using 5 mM GTP. E483K, E576K and L618V displayed GTP concentration-dependent release of pyrophosphate, resulting in saturation Michaelis-Menten kinetics (Tab. 2). **d.** PDE enzymatic kinetics determined by MANT-c-di-GMP hydrolysis. MANT-c-di-GMP hydrolysis curves for wild type, E483K, E576K and L618V (Tab. 3). Error bars represent standard error of the mean. Kinetics were performed in two independent measurements.

**Table 2.** Kinetic parameters for DGC activity of MbaA and evolved mutants.

|  | Wild-type<sup>a</sup> | E483K<sup>a</sup> | E576K<sup>a</sup> | L618V<sup>a</sup> |
| --- | --- | --- | --- | --- |
| $k_{\text{cat}}$ (s<sup>-1</sup>) | < 0.002 | 0.086 ± 0.013 | 0.037 ± 0.013 | 0.023 ± 0.003 |
| $K_M$ (μM) | NC | >5000 | 2500 ± 1500 | 600 ± 250 |
| $k_{\text{cat}}/K_M$ (M<sup>-1</sup> s<sup>-1</sup>) | NC | 9 ± 1 | 14 ± 4 | 40 ± 10 |

<sup>a</sup> Determined at 30°C by determining the release of P<sub>i</sub> during the production of c-di-GMP from GTP.
NC: not calculated due to lack of activity.
Errors are reported as the standard error of the mean.

**Table 3.** Kinetic parameters for PDE activity of MbaA and evolved mutants.

|  | Wild-type<sup>a</sup> | E483K<sup>a</sup> | E576K<sup>a</sup> | L618V<sup>a</sup> | E553A<sup>a</sup> |
| --- | --- | --- | --- | --- | --- |
| $k_{\text{cat}}$ (s<sup>-1</sup>) | 0.026 ± 0.003 | 0.002 ± 0.001 | 0.011 ± 0.005 | 0.001 ± 0.0002 | -<sup>b</sup> |
| $K_M$ (μM) | 11 ± 3 | 13 ± 6 | 16 ± 11 | 6 ± 3 | -<sup>b</sup> |
| $k_{\text{cat}}/K_M$ (M<sup>-1</sup> s<sup>-1</sup>) | 2,000 ± 500 | 150 ± 40 | 700 ± 200 | 250 ± 70 | 380 ± 30 |

<sup>a</sup> Determined at 37°C using MANT-c-d-GMP as a substrate.
<sup>b</sup> Could not be calculated due to the high degree of linearity (non-saturation kinetics).
Errors are reported as the standard error of the mean.

To investigate how mutations in the GGDEF (E483K) and EAL domain (E576K, L618V) affect the activity of MbaA, we purified the cytoplasmic part of wild-type MbaA and mutant variants with a MBP tag and determined steady-state enzyme kinetics. To measure their DGC activity, we monitored the release of pyrophosphate (Pi) using a coupled enzymatic assay^46^. In this assay, Pi, which is released during the formation of c-di-GMP from GTP, is measured by monitoring the reaction of the released Pi with 2-amino-6-mercapto-7-methlpurine (MESG) and serves as an indirect measure of DGC activity (Fig. 3c. and Table 2, Fig. S4a)^46–48^. The incubation of wild-type MbaA with 5 mM GTP did not result in the release of Pi concentration above the background reaction (Fig. 3c), supporting the notion that wild-type MbaA predominantly hydrolyzes c-di-GMP^22,25^. In contrast, E483K, E576K, and L618V generated a substantial release of Pi. The release of Pi followed saturation kinetics with a *k*cat/*K*M ranging from 9 - 40 s^-1^ M^-1^. E483K, E576K, and L618V exhibited *K*M values ranging from 600 to >5000 µM and *k*cat values 11.5 to 43-fold above the background reaction (Table 2). The reaction velocity of pure Pi with MESG (Fig. S4b) was several magnitudes higher than the determined *k*cat values obtained during GTP turnover, demonstrating that the determined *k*cat values mainly represent the formation of c-di-GMP and not the secondary reaction (Fig. 3, Fig. S4b and Table 2). Consequently, mutations in both the GGDEF domain (E483K) and the EAL domain (E576K, L618V) increase the DGC activity of MbaA *in vitro*.

Next, we employed fluorescence spectroscopy to determine the effect of the mutations on the PDE activity of MbaA using MANT-c-di-GMP (Fig. 3d and Table 3)^49^. To validate our assay, we tested MANT-c-di-GMP hydrolysis by wild-type MbaA and a mutant with an active site mutation in the EAL domain (E553A). Wild-type MbaA hydrolyzed MANT-c-di-GMP following a saturation kinetic (*K*M = 11 µM, *k*cat = 0.026 s^-1^, and *k*cat/*K*M = 2,000 M^-1^ s^-1^). In contrast, the E553A mutant had a 5-fold lower *k*cat/*K*M and non-saturation kinetics (linear), demonstrating that E553A disrupts PDE activity, potentially by impairing substrate affinity (Fig. S4d). After confirming that our system allows quantifying MANT-c-di-GMP hydrolysis, we determined the PDE activity of the E483K, E576K, and L618V mutants (Fig. 3d, Table 3). All mutations substantially decreased *k*cat/*K*M by 3 to 13-fold compared to wild type MbaA. Like wild type MbaA, the GGDEF (E483K) and EAL domain (E576K and L618V) mutants followed saturation kinetics with overall similar substrate affinities (*K*M ranging from 6 to 16 µM). In contrast, E483K, E576K, and L618V mutants each exhibited a reduction in turnover compared to the wild type, with a 13-, 2-, and 26-fold decrease in *k*cat, respectively (Tab. 3).

Our kinetic investigation aligns well with our *in vivo* phenotypic results and demonstrate that the mutations in both the GGDEF (E483K) and EAL domain (E576K and L618V) activate MbaA’s DGC activity, leading to increased synthesis of c-di-GMP, while simultaneously compromising the PDE activity, reducing hydrolysis of c-di-GMP. Together, these results suggest that the hyper-biofilm phenotype selected for during biofilm evolution is seemingly a result of simultaneously modulating both the DGC and PDE activity of MbaA.

### Mutations in MbaA decouple the enzyme from its polyamine-mediated regulation

Post-translational regulation via polyamines like spermidine and norspermidine modulates the enzymatic activity of MbaA through the periplasmic solute-binding protein, NspS^22,23,25,28,30,31^. Norspermidine acts as a biofilm stimulator that increases the DGC activity of MbaA, while spermidine inhibits biofilm formation and stimulates the PDE activity of MbaA^22,23,25,30,31^. Next, we aimed to investigate how these mutations impact the post-translational regulation of MbaA.

First, we investigated the effect of norspermidine and spermidine on biofilm formation in *V. cholerae* C6706 producing wild-type MbaA. In agreement with previous studies^22,23,25,31^, the addition of 1 mM norspermidine, increased biofilm formation 10-fold compared to cultures grown in LB alone (Fig. 4a, paired one-way ANOVA, *P* <0.0001) and led to formation of wrinkly biofilm pellicles (Fig. 4c). In contrast, the addition of 2 mM spermidine significantly reduced biofilm formation by 1.6-fold (Fig. 4a, paired one-way ANOVA, *P* = 0.0006).

**Figure 4:**
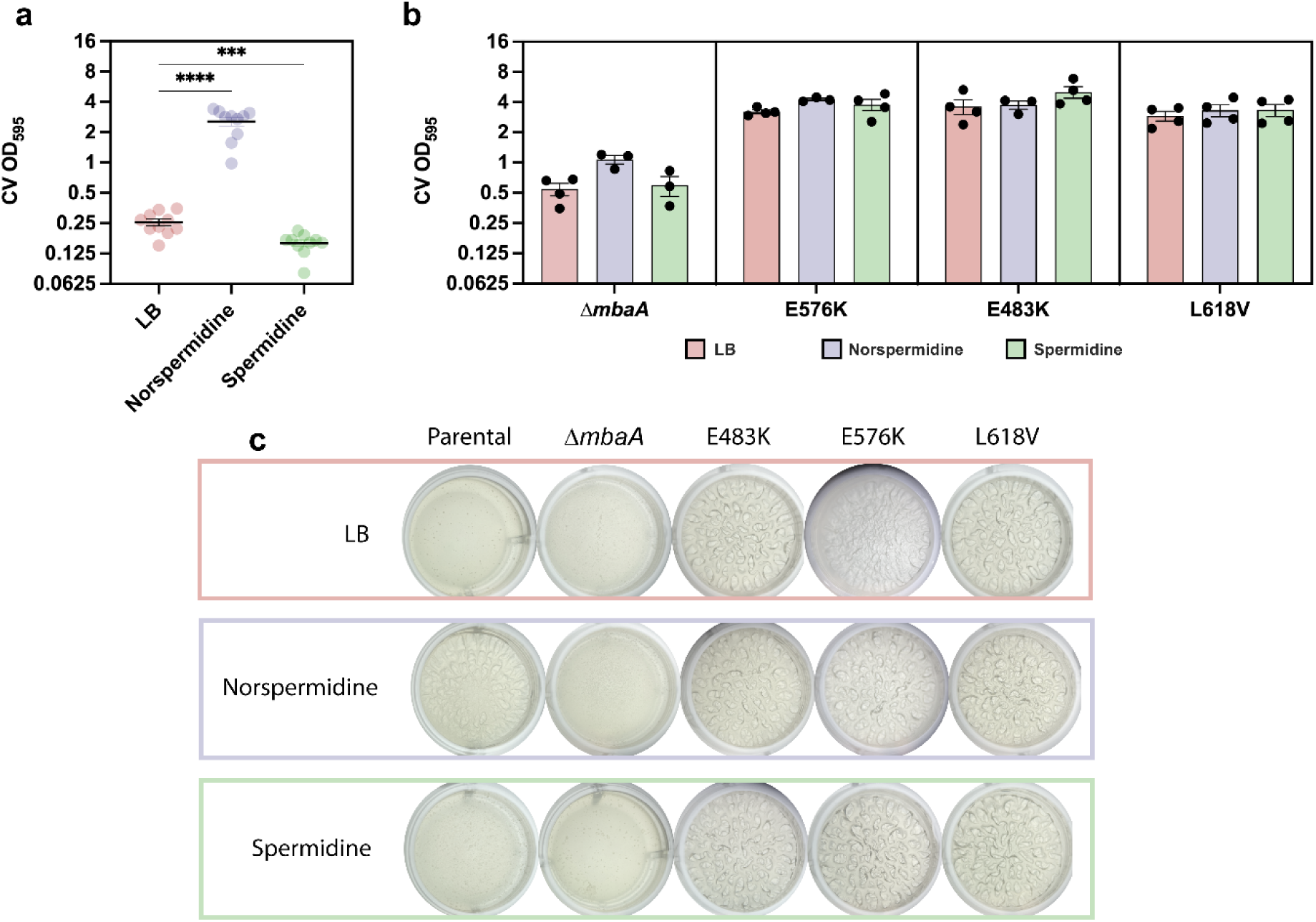
Evolved and identified mutations desensitize MbaA to polyamines. **a.** Biofilm formation of *V. cholerae* C6706 producing wild-type MbaA in LB or LB supplemented with either norspermidine or spermidine. Statistical significance was tested with a paired one-way ANOVA followed by Dunnett’s multiple comparisons tests, ***; *P* < 0.001, ****; *P* <0.0001. **b.** Biofilm formation of *V. cholerae* C6706 carrying MbaA variants (Δ*mbaA*, E576K, E483K and L618V) in LB or LB supplemented with either norspermidine or spermidine. **c.** Representative images of morphology of biofilm pellicles formed by either the parental *V. cholerae* C6706, or strains carrying either Δ*mbaA*, E576K, E483K or L618V in LB or LB supplemented with either norspermidine or spermidine. For panels **a** and **b**, biofilm production was quantified by measuring the absorbance at 595 nm of dissolved biofilms stained with crystal violet (CV). Bar represents means and each dot represents one biological replicate. Error bars represent the standard error of the mean.

Next, we tested the ability of a *mbaA* deletion mutant (Δ*mbaA*) to produce biofilms in the presence and absence of norspermidine or spermidine (Fig. 4b). The Δ*mbaA* mutant did not respond to either norspermidine or spermidine excluding the presence of other polyamine responsive biofilm regulators in *V. cholerae* (Fig. 4b-c).

Finally, the behavior of *V. cholerae* producing either the E483K, E576K, or L618V mutants was determined. Neither norspermidine nor spermidine affected the strains’ ability to produce biofilms (Fig. 4b). Furthermore, all mutants produced wrinkly pellicles in the presence of both polyamines (Fig. 4c). These findings demonstrate that the biofilm-evolved mutations in the GGDEF (E483K) and EAL domain (E576K, L618V) likely decouple MbaA from polyamine-mediated regulation via NspS, seemingly locking it into a constitutively active DGC state (Fig. 4b-c).

## DISCUSSION

As a key regulator of bacterial behavior, genetic changes in the c-di-GMP signaling system represent a potent strategy to trigger *in vivo* and *in vitro* adaptation across various environments^6,11–16,50–56^. Here, during experimental evolution of *V. cholerae* biofilm pellicles, we demonstrate strong selection for a biofilm-associated lifestyle through mutations in the bifunctional c-di-GMP-metabolizing enzyme MbaA (Fig. 1d-e). Our findings are consistent with the current paradigm, stating that in its native state MbaA hydrolyzes c-di-GMP^22,23,25,28,31^. We showcase that biofilm evolution can select for mutations in MbaA that alter its enzymatic activity and decouple the enzyme from its post-translational polyamine regulation. Ultimately, these changes likely lead to constitutive production of c-di-GMP, thereby inducing a switch in lifestyle from planktonic to biofilm-associated (Tab. 1-3, Fig. 2-5, Fig. S3).

**Figure 5:**
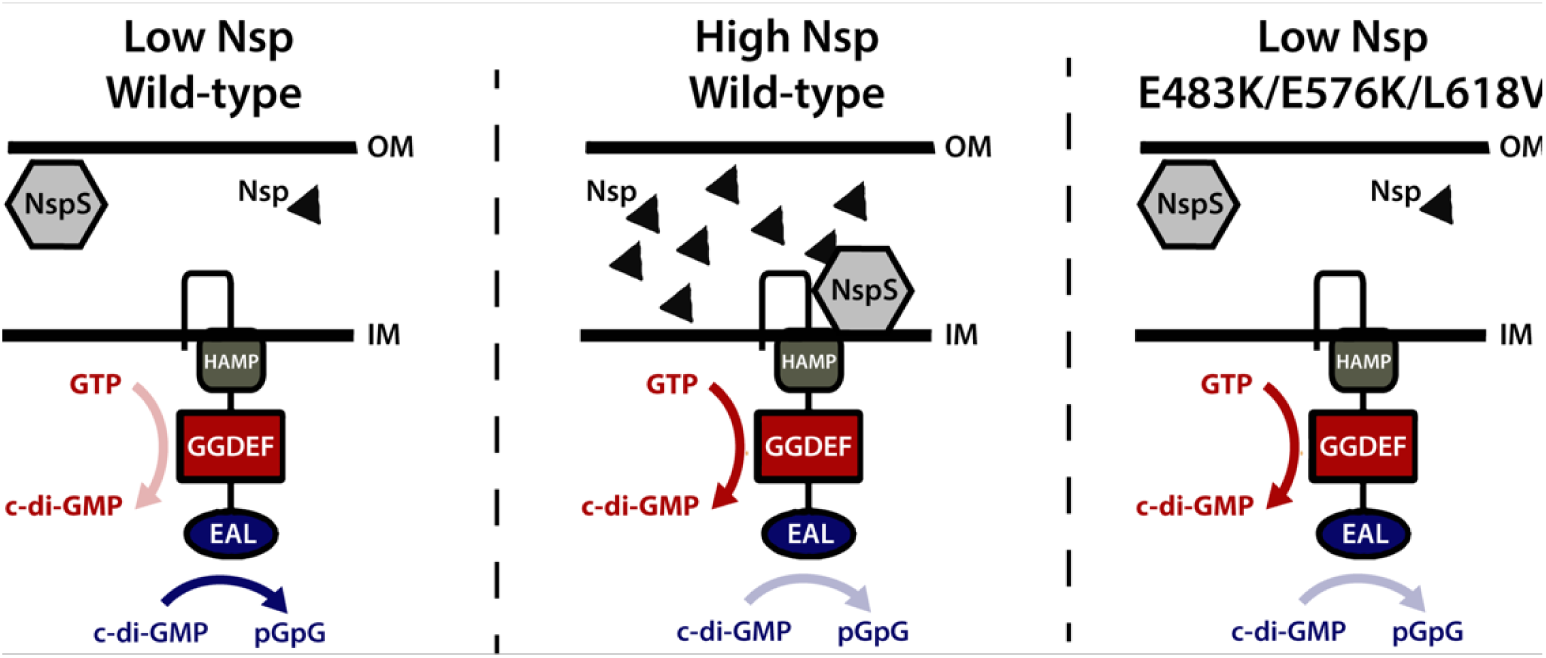
Mutations in MbaA can lead to constitutive production of c-di-GMP. Model of wild-type membrane-bound MbaA and the periplasmic polyamine sensor NspS, which is located between the outer (OM) and inner membrane (IM) as proposed previously^22^ followed by model of the effect of the selected mutations (E576K, E483K and L618V). E576K, E483K and L618V mutations in MbaA activate the DGC activity of the GGDEF domain and reduce the PDE activity of the EAL domain, thereby inducing c-di-GMP-synthesis even in the absence of norspermidine (Nsp) or despite the presence of spermidine. Figure is adapted from work by Bridges et al^22^.

Our findings show that mutations in the EAL domain (E576K and L618V) increase the DGC activity of the GGDEF domain, suggesting that interdomain interactions regulate the enzymatic activity of MbaA (Fig. 2,3 and 5, Tab. 1). In support of this, Bridges et al.^22^ demonstrated that the removal of either enzyme domain in MbaA impairs the enzymatic activity of the remaining one. Furthermore, mutations in the EAL domain of the bifunctional c-di-GMP-metabolizing enzyme DcpA from *Mycobacterium smegmatis* negatively affected both DGC and PDE activity^57,58^. Additionally, a study in *Pseudomonas fluorescens* showcased that mutations in the EAL domain (E1081A and E1082K which correspond to E576 and E577 in MbaA) of the bifunctional c-di-GMP-metabolizing enzyme MorA induced a phenotype associated with increased levels of c-di-GMP, suggesting that such mutations can stimulate DGC activity^55^. The GGDEF and EAL domains of c-di-GMP-metabolizing enzymes have previously been shown to interact *via* electrostatic interactions^44,45^. The fact that disturbing these electrostatic interactions, as demonstrated in MorA, can induce phenotypes associated with altered levels of c-di-GMP suggests that interdomain crosstalk modulates the overall enzymatic activity^44,45,55^. We speculate that such crosstalk may be involved in modulating the enzymatic activity of the here evolved MbaA variants. However, further structural biology studies are required to confirm this.

Our data demonstrate that biofilm evolution selects for mutations that decouple MbaA from polyamine-mediated regulation *via* NspS (Fig. 4b-c, 5). Many c-di-GMP-metabolizing enzymes are under strict post-translational regulation^17–21^. This ensures tight regulation of the cellular pool of c-di-GMP but also represents an adaptive target in bacteria. Prior studies have demonstrated that mutations in regulatory proteins or sensory domains of c-di-GMP-metabolizing enzymes can shift the intracellular c-di-GMP equilibrium, thereby inducing adaptive changes^6,11,14,15,17,18,50,52–55^. C-di-GMP is a key modulator of bacterial behavior, not solely biofilm regulation, and has been demonstrated to affect other bacterial traits (e.g., motility, virulence, and cooperative behavior)^17,18,50,52,53,55,59^. Decoupling of c-di-GMP- metabolizers from their regulators has been described in multiple enzymatic systems, such as Wsp and YfiBNR^12,14,50–53,60^. Our findings provide additional support for the emerging realization that regulatory decoupling of c-di-GMP-metabolizing enzymes may be a widespread and important evolutionary mechanism for bacterial adaptation.

In conclusion, our study demonstrates how biofilm evolution selects for mutations in the bifunctional c-di-GMP-metabolizing enzyme MbaA, triggering the transition to a biofilm- associated lifestyle in *V. cholerae*. These mutations activate the DGC activity of the GGDEF domain while reducing the PDE activity of the EAL domain and decouple MbaA from polyamine-mediated regulation by NspS. Consequently, this study sheds light on how rewiring of existing c-di-GMP metabolizing enzymes can induce lifestyle transitions in pathogens. A detailed molecular understanding of the evolution and regulation of the c-di-GMP signaling system, a conserved regulator of bacterial behavior, can aid the development of targeted interventions against biofilm-associated infections.

## MATERIAL AND METHODS

### Strains and media

Strains and oligonucleotides used in this study are listed in Table S1-2. Unless otherwise stated, cultures were grown in premixed Lysogeny-Broth-Miller (LB) media (10 g/L of tryptone, 10 g/L of sodium chloride and 5 g/L of yeast extract) (Sigma-Aldrich). Growth media were supplemented with streptomycin 100 μg/ml (Sigma-Aldrich), carbenicillin 50 μg/ml (Sigma-Aldrich), 100 μg/ml ampicillin (Sigma-Aldrich), spermidine 2 mM (Sigma-Aldrich) and norspermidine 1 mM (Sigma-Aldrich) as indicated.

### Biofilm pellicle evolution

*V. cholerae* C6706 was grown on LB agar plates and a single colony for each of the eight populations was selected and used to inoculate a 2 ml LB liquid culture in a 24-well plate (Corning, USA), which was grown overnight at 37 °C with shaking (225 rpm) in a Multitron incubation shaker (Infors HT). The following day the overnight culture was diluted 1:100 into 2 ml of fresh LB-medium in a 24-well plate (Corning, USA) and grown statically for 24 hours at 30 °C. After 24 hours, all the bacterial cultures contained biofilm pellicles at the air-liquid interface. These biofilm pellicles were harvested with a sterile loop and transferred into 1 ml of sterile PBS (Sigma-Aldrich, 37 mM NaCl, 2.7 mM KCl and 10 mM phosphate buffer solution, pH 7.4). The harvested biofilm pellicle was broken down by vortexing continuously for 60 seconds. After vortexing, the bacterial PBS-suspension was filtered through a Whatman® Puradisc 5 μm (Sigma-Aldrich) sterile filter to remove larger biofilm aggregates and isolate biofilm-derived bacterial cells. The bacterial filtrate was used to inoculate a new static culture as described above. This process was repeated every 24 hours for a total of eight days. The pellicle evolution was performed in six parallel lineages. For propagation of the planktonic populations (*n*=2), the biofilm pellicles were harvested and discarded, and the planktonic cultures were transferred at 1:100 ratio into fresh LB medium.

### Strain and plasmid construction

All chromosomal genetic modifications (whole gene knockouts, partial gene knockouts or point mutations) were created by allelic exchange^61^. To this end, 1000 to 1200 base pairs of homologous regions upstream and downstream of the locus of the desired genetic change were PCR-amplified (Phusion polymerase from New England Biolabs) using primers (listed in Table S2). All oligonucleotides were synthetized by Sigma Aldrich and purified to remove salts. The homologous regions were integrated into the suicide plasmid pCVD442 digested by *XbaI* through a Gibson assembly reaction to create a recombinant allelic-exchange vector^62,63^. *E. coli* DH5αpir was used as host bacterium to amplify the recombinant vectors^64^. The correct construction of the recombinant allelic-exchange vector was confirmed by Sanger sequencing (primers see Table S2) the insert of the pCVD442-based recombinant suicide vectors (Genewiz). After confirmation that the recombinant allelic-exchange vector contained the correct insert, it was isolated with a miniprep kit (Qiagen) and chemically transformed into *E. coli* SM10λpir as previously described^65,66^.

To generate strains with chromosomal modifications, the pCVD442-based recombinant allelic-exchange vectors described above were transferred into *V. cholerae* C6706 by conjugation. Afterwards, transconjugants were isolated on LB-plates with streptomycin and carbenicillin to select for transconjugants with a chromosomally integrated vector (single crossover). Transconjugants with the appropriate resistance pattern were then grown on 10% sucrose plates to select for double crossovers. Bacterial clones growing on sucrose were screened by streaking them on two different LB plates supplemented with either streptomycin (100 μg/ml) or carbenicillin (50 μg/ml). Strains that only grew on LB supplemented with streptomycin were verified by PCR with primers (Table S2) that bound outside of the mutated locus. Finally, to confirm that the desired genetic change was present, genomic DNA of isolated clones was isolated, and Sanger sequenced (Genewiz).

### Biofilm formation assay

Biofilm formation was determined using crystal violet staining as previously described^67^. Briefly, overnight cultures of single colonies were grown in LB medium at 37°C with shaking (225 rpm). 2 ml LB medium was inoculated 1:100 in 14 ml culture tubes and incubated for 12 hours with shaking at 225 rpm. After 12 hours, the media was removed from the culture tubes and the tubes were gently submerged in distilled water three times to remove non-adherent cells. Afterwards, the attached biomass was left to dry at room temperature. To stain the attached biomass, 3 ml 0.1% crystal violet (Sigma-Aldrich) was added to each tube and incubated for 10 minutes. Afterwards, the 0.1% crystal violet solution was removed, and the tubes were gently washed three times in distilled water to remove residual crystal violet. Stained biomass was left to dry at room temperature and the retained dye was subsequently dissolved in 3 ml 30% acetic acid. Finally, biomass was quantified by measuring the absorbance of the dissolved dye at 595 nm in an Epoch2 plate reader (Biotek). For measurement of biofilm formation in the presence of polyamines (spermidine or norspermidine), the assay was performed in LB supplemented with either norspermidine 1 mM or spermidine 2 mM. Furthermore, to allow higher levels of biofilm formation with the goal of increasing the detection limit of biofilm inhibition, cultures were grown for 24 hours with shaking instead of 12 hours.

### Assessment biofilm pellicle morphology

Cultures were prepared from frozen stocks in 14 mL culture tubes containing 2 mL LB medium and incubated overnight at 37 °C with shaking (225 rpm). The following day, overnight cultures were diluted 1:100 into 2 ml of fresh LB medium in a 24-well plate (Corning, USA) and incubated statically at 37°C for 24 hours. Pellicle formation was visualized and imaged with a Euromex NexiusZoom (Euromex, Netherlands) stereo microscope at 6.7x magnification after 24 hours.

### Whole-Genome Sequencing

For short-read illumina whole genome sequencing, genomic DNA from the eight evolved populations (evolved populations 1-8) was isolated with a GenElute Bacterial Genomic DNA Kit (Sigma Aldrich). Purity and concentration of gDNA was assessed with a NanoDrop ND- 1000 spectrophotometer (Thermo Scientific) and a Qubit Fluorometer (Thermo Scientific). Short-read sequencing library preparation were performed following the manufacturers’ instructions and the samples were sequencing at the Genomic Support Centre Tromsø, UiT the Arctic University of Norway. The Nextera XT DNA Library preparation kit (Illumina) was used with an input of 1 ng genomic DNA and dual indexes. Samples were sequenced on a NextSeq 550 instrument (Illumina) with 300 cycles (2 × 150 bp paired end reads). A single high-output flow cell was used for the eight population samples aiming to achieve 1000x coverage.

### Short-Read Sequence Analysis

The BreSeq computational pipeline v0.33.0 was used to predict mutations in the evolved population samples using short-read sequencing data^68^. Sequence reads of all evolved populations were mapped against the reference genome of our previously sequenced stock of *V. cholerae* C6706 (accession number: CP157384, CP157385)^69,70^. BreSeq was run with default settings in full polymorphism mode for population samples with the following exceptions: Consensus mode (Frequency-cutoff 0.9, minimum-variant-coverage 10, consensus-minimum-total-coverage 10), polymorphism-mode (frequency-cutoff 0.01, minimum-variant-coverage 10, minimum-total-coverage 100, base quality score 20). In addition, short-read sequencing data from our frozen stock of parental *V. cholerae* C6706 was used as input in BreSeq and compared against the C6706 genome (accession number: CP157384, CP157385)^70,71^. Mutations predicted by BreSeq in the evolved populations were excluded from the analysis when they were also predicted in the short-read sequencing data from the parental *V. cholerae* C6706. Due to the limitation of short-read sequencing technology only SNPS and small InDels were analyzed and chromosomal inversions, large rearrangements, and mutations in repeat regions were excluded.

### Multiple sequence alignments

Multiple sequence alignments were conducted with the ENDscript server using GGDEF-EAL domain proteins with GGDEF and EAL domains from *V. cholerae* C6706 and well characterized GGDEF and EAL domains from other bacterial species^72^. MorA (PDB ID: 4RNH) was used as reference for the protein 3D structure^45^.

### Homology modelling

To obtain a homology modelled structure of MbaA we employed SWISS-MODEL and the solved structure of MorA (PDB ID: 4RNH) with sequence identity of 35.2% as template^45,73^.

### Protein expression and purification

To purify MbaA, we created a pDEST17-based expression vector encoding non-membrane-bound wild-type MbaA or mutant variants (GGDEF-EAL domain, L328 – R791) linked to a maltose-binding protein (MBP). The vector was created by PCR-amplifying (Phusion polymerase from New England Biolabs) wild-type MbaA from genomic DNA extracted from *V. cholerae* C6706. Afterwards, the amplified DNA encoding MbaA was cloned onto the pDEST17-based vector through a Gibson assembly reaction^62^. The pDEST17-based expression vector containing non-membrane-bound wild-type MbaA was used as template for site-directed mutagenesis to create the mutant variants of MbaA. The final constructs were confirmed by Sanger sequencing (GeneWiz).

The protein was expressed in and purified from *E. coli* BL21 AI^74^. Cultures were grown in 1 L LB supplemented with 0.2% glucose and 100 mg/L ampicillin at 37°C. Expression of the MBP-tagged enzymes was induced when cultures reached an optical density (OD600) of approximately 0.5 by adding L-arabinose to a final concentration of 0.2%. Expression was performed for 16 hours at 30°C and cultures were harvested by centrifugation for 30 min at 4000 x g at 4 °C. Cells were resuspended in the column buffer (200 mM NaCl, 0.5 mM EDTA, 5 mM MgCl2, 20 mM Tris-HCl, pH 7.6) with 1x proteinase inhibitor (Thermo Scientific) and sonicated for 30 min on ice. Cell debris were separated by centrifugation at 4°C, 20,000 x g for 1 h and supernatant incubated with 1 mL amylose. After 16 hours incubation at 4°C, amylose was washed with 20 mL column buffer and protein elution was performed stepwise by adding column buffer supplemented with 20 mM L-Maltose. Maltose was removed using molecular cut off filters (50,000 Da, Merck). Proteins extinction coefficient was determined using ProtParam and the protein concentration was determined using Nanodrop at OD ^75^.

### Steady state enzyme kinetics for DGC activity

The formation of c-di-GMP was determined by using the EnzCheck pyrophosphate kit (Invitrogen), by measuring the Pi that is released during the cyclization of GTP. Pi and MESG are converted by the purine nucleoside phosphorylase (PNP) enzyme to ribose 1-phosphate and 2-amino-6-mercapto-7-methylpurine which causes a shift in absorbance maximum from 330 nm to 360 nm^46–48^. We determined the extinction coefficient for the reaction of Pi with MESG in 50 mM Tris HCl pH 7.6 supplemented with 10 mM MgCl2, 0.5 mM EDTA and 50 mM NaCl at 30°C to be 18,000 [M^-1^ cm^-1^] (Fig. S4a). The reaction product was monitored at 360 nm (Epoch 2, Biotek). Using the determined extinction coefficient, we performed Michaelis Menten kinetics on wild-type MbaA and the corresponding mutants (1 μM final protein concentration). GTP was preincubated with all buffer ingredients for 10 min at room temperature before adding MbaA or the corresponding mutants. GTP concentrations ranging from 0.25 to 5 mM to determine the kinetic parameters *k*cat, *K*M and *k*cat/*K*M by fitting the curves to Eq. 1 (velocity = Vmax*(S)/ (*K*M + (S))) in GraphPad Prism v.10.2.3. Analyses were performed at least in duplicates in 96 well plates (Nunc) and at a final volume of 100 μL.

### Steady state enzyme kinetics for PDE activity

Phosphodiesterase activity of wild-type MbaA and mutants was determined using a final enzyme concentration of 2 μM enzyme and 50 mM Tris-HCl pH 8.0 (Sigma-Aldrich) buffer, supplemented with 5 mM MnCl2 (Sigma-Aldrich). Catalytic activity was determined at 37 °C using the fluorescent (λexc = 355 nm) c-di-GMP derivate 2’-O-(N’-methylanthraniloyl)-cyclic diguanylate (MANT-c-di-GMP) and changes in fluorescence were monitored at λemi = 448 nm (Gain = 97; excitation band with 20 nm; Spark, Tecan)^76^. Extinction coefficient for the hydrolysis of MANT-c-di-GMP was determined by analyzing emitted fluorescence of MANT-c-di-GMP before and after incubation with 2 μM wild-type MbaA for 2 hours (Fig. S4c). The change in fluorescence was used to calculate the extinction coefficient of -2,850 [μM^-1^ cm^-1^]. Reaction curves were fitted to Eq 1. (GraphPad Prism v.10.2.3) and *k*cat, *K*M and *k*cat/*K*M were calculated based on the extinction coefficient in a final volume of 50 μL.

### Statistical analysis

The normality of the datasets was investigated by the Kolmogorov-Smirnov test and by visual inspection of the residual plots. Statistical significance was tested with one-way ANOVA followed by either Dunnett’s T3 or Dunnett’s multiple comparisons tests. Statistical significance was set at α < 0.05. All statistical analyses were performed using GraphPad Prism v.10.2.3.

## FUNDING, ACKNOWLEDGMENT AND CONFLICT OF INTERESTS

The authors declare no conflict of interest. CF was supported by the Centre for New Antibacterial Strategies (CANS) at UiT – The Arctic University of Norway and the and Northern Norway Regional Health Authority (HNF1722-24). ØML was funded by UiT – The Arctic University of Norway. PJJ thanks The Olav Thon Foundation for funding. SA thanks the Norwegian Research Council (NFR) 249979 for funding. The funders had no role in study design, data collection and interpretation, or the decision to submit the work for publication. We thank Julia Maria Kloos and João Alves Gama for their assistance with identifying mutations in the evolved populations with BreSeq. We also thank the Genomic Support Centre Tromsø at UiT – The Arctic University of Norway for conducting read sequencing library preparation and whole genome sequencing of evolved populations.

## AUTHOR CONTRIBUTION

ØML conceived the study. ØML and CF performed the experimental work. All authors interpreted experimental results. ØML and CF wrote the original manuscript draft, all authors edited and revised the manuscript.

## DATA AVAILABILITY STATEMENT

The data underlying the figures in this article are published alongside the article as source file. Sequencing data generated in this study will be made available in a public database for sequencing data (NCBI).

