## Supplemental materials for "Biofilm selection constitutively activates c-di-GMP synthesis by the bifunctional enzyme MbaA"

#### Contents

1. **Supplementary tables**

**Table S1.** Strains constructed and used in this study.

**Table S2.** Primers used in this study

**Table S3.** Biofilm formation of wild-type and evolved populations of V. cholerae

**Table S4.** P-values from one-way ANOVA comparing biofilm formation of all mbaA variants

2. **Supplementary figures**

**Figure S1.** Alignment of selected EAL-domains from GGDEF-EAL proteins

**Figure S2.** Alignment of selected GGDEF-domains from GGDEF-EAL proteins

**Figure S3.** Biofilm formation of parental *V. cholerae* and all tested MbaA mutants

**Figure S4.** Validation of indirect measurements of DGC- and PDE-activity

3. **References**

### 1. Supplementary tables

**Table S1: Strains constructed and used in this study.**

| Strain no. | Strain background [a] | Plasmid | mbaA genotype | Description | Reference |
|---|---|---|---|---|---|
| SA43-77 | C6706 | None | Wild-type | Parental strain, quorum sensing deficient | [1,2] |
| SA 44-45 | C6706 | None | - | Planktonic-evolved population 1 | This study |
| SA 44-49 | C6706 | None | - | Planktonic-evolved population 2 | This study |
| SA 44-42 | C6706 | None | - | Biofilm-evolved population 1 | This study |
| SA 44-43 | C6706 | None | - | Biofilm-evolved population 2 | This study |
| SA 44-44 | C6706 | None | - | Biofilm-evolved population 3 | This study |
| SA 44-46 | C6706 | None | - | Biofilm-evolved population 4 | This study |
| SA 44-47 | C6706 | None | - | Biofilm-evolved population 5 | This study |
| SA 44-48 | C6706 | None | - | Biofilm-evolved population 6 | This study |
| SA 45-35 | C6706 | None | ∆mbaA | - | This study |
| SA 47-5 | C6706 | None | ∆EAL | - | This study |
| SA 48-56 | C6706 | None | E483K | - | This study |
| SA 47-2 | C6706 | None | G425A | - | This study |
| SA 47-4 | C6706 | None | G425A/E576K | - | This study |
| SA 48-78 | C6706 | None | E553A | - | This study |
| SA 46-46 | C6706 | None | E576K | - | This study |
| SA 47-72 | C6706 | None | L618V | - | This study |
| SA 48-51 | *E. coli* BL21 AI | pOML213 | Wild-type | Expression vector | This study |
| MP32-46 | *E. coli* BL21 AI | pOML213 | E483K | Expression vector | This study |
| SA 48-52 | *E. coli* BL21 AI | pOML213 | E576K | Expression vector | This study |
| MP-32-42 | *E. coli* BL21 AI | pOML213 | L618V | Expression vector | This study |
| MP-32-48 | *E. coli* BL21 AI | pOML213 | E553A | Expression vector | This study |

**Cloning strains:**

| Strain no. | Strain background a | Plasmid | Insert | Reference |
|---|---|---|---|---|
| SA 36-9 | *E. coli* DH5αpir | None | None | [3] |
| SA 22-7 | *E. coli* SM10λpir | None | None | [4] |
| SA 48-30 | *E. coli* DH5αpir | pCVD442 | None | [5] |
| SA 45-23 | *E. coli* DH5αpir | pOML194 | *∆mbaA* | This study |
| SA 45-28 | *E. coli* SM10λpir | pOML194 | *∆mbaA* | This study |
| SA 46-20 | *E. coli* DH5αpir | pOML195 | mbaA_E576K | This study |
| SA 46-79 | *E. coli* SM10λpir | pOML195 | mbaA_E576K | This study |
| SA 46-68 | *E. coli* DH5αpir | pOML197 | mbaA_G425A | This study |
| SA 46-77 | *E. coli* SM10λpir | pOML197 | mbaA_G425A | This study |
| SA 46-67 | *E. coli* DH5αpir | pOML198 | mbaA_∆EAL | This study |
| SA 46-76 | *E. coli* SM10λpir | pOML198 | mbaA_∆EAL | This study |
| SA 48-31 | *E. coli* DH5αpir | pOML216 | mbaA_E483K | This study |
| SA 48-32 | *E. coli* SM10λpir | pOML216 | mbaA_E483K | This study |
| SA 48-70 | *E. coli* DH5αpir | pOML222 | mbaA_E553A | This study |
| SA 48-71 | *E. coli* SM10λpir | pOML222 | mbaA_E553A | This study |
| MP32-43 | *E. coli* E.cloni | pOML213 | mbaA_E483K | This study |
| SA48-44 | *E. coli* DH5αpir | pOML213 | mbaA_E576K | This study |
| SA48-76 | *E. coli* DH5αpir | pOML213 | mbaA_L618V | This study |
| MP32-45 | *E. coli* E.cloni | pOML213 | mbaA_E553A | This study |

**Table S2: Primers used in this study**

| No. | Name | Sequence (5’ to 3’) |
|---|---|---|
| 1409 | P1409_VC0703_KO_homol_1_for | aggtatatgtgatgggttaaaaaggatcgatcctaacgctttctgtcgctgatg |
| 1410 | P1410_VC0703_KO_homol_1_rev | ttatgattggaggcatgaagccatggggagatctccaccggtgcaatcagtaaca |
| 1411 | P1411_VC0703_KO_homol_2_for | taaaccatagaattctgttactgattgcaccggtggagatctccccatggcttca |
| 1412 | P1412_VC0703_KO_homol_2_rev | ccgggagagctcgatatcgcatgcggtacctctagatttgatgggcgtacaaggc |
| 1413 | P1413_VC0703_KO_seq | cggtctcttacagtggcgat |
| 1414 | P1414_VC0703-KO_PCR_for | tcgctggtttcgaccttact |
| 1415 | P1415_VC0703-KO_PCR_rev | gaccagaaccgaatcttggc |
| 1490 | P1490_VC0703_EAL_KO_ho-mol_1_for | aggtatatgtgatgggttaaaaaggatcgatccttctcaacgccgctcagttat |
| 1491 | P1491_VC0703_EAL_KO_homol_1_rev | cggcattcactttggctgggttctgtgaagcaatacgcctgcgaatagtaagcgac |
| 1492 | P1492_VC0703_EAL_KO_ho-mol_2_for | cgggcaaaaaccaagtcgcttactattcgcaggcgtattgcttcacagaacccagc |
| 1497 | P1497_VC0703_do-main_KO_seq_F | ccacccacaaaccgacaatc |
| 1498 | P1498_VC0703_do-main_KO_seq_R | tcaactgcgttttcctgtcg |
| 1503 | P1503_G1274C_VC0703_homol_1_for | aggtatatgtgatgggttaaaaaggatcgatcctcagttatgccttcacgctgg |
| 1504 | P1504_G1274C_VC0703_homol_1_rev | gtttgggggagacgagcaaaatagcaaactcatcagcagataaacgggccgcaata |
| 1505 | P1505_G1274C_VC0703_homol_2_for | ccataacacctacagtattgcggcccgtttatctgctgatgagtttgctattttgctcg |
| 1506 | P1506_G1274C_VC0703_homol_2_rev | ccgggagagctcgatatcgcatgcggtacctctaggcattcactttggctgggtt |
| 1595 | P1595_mbaA_G2464A_homol_1_F | aggtatatgtgatgggttaaaaaggatcgatcctccgatccatttgccgaactt |
| 1596 | P1596_mbaA_G2464A_homol_1_R | atacatggcggtatcggcattgagcagcaatttcttaatgtgttcgccatctttaggg |
| 1597 | P1597_mbaA_G2464A_homol_2_F | gtattgcgaccttccctaaagatggcgaacacattaagaaattgctgctcaatgccg |
| 1598 | P1598_mbaA_G2464A_homol_2_R | ccgggagagctcgatatcgcatgcggtacctctaggctgtttggcatttcggact |
| 1669 | P1669_mbaA_E553A_homol1_F | aggtatatgtgatgggttaaaaaggatcgatcctggtggtgtatgtggtgcttg |
| 1670 | P1670_mbaA_E553A_homol1_R | caccgagcaattcagattgccagcgcagcaatactgcgaagcccactaagcgtttcc |
| 1671 | P1671_mbaA_E553A_homol2_F | ctttacttgctcagggaaacgcttagtgggcttcgcagtattgctgcgctggcaat |
| 1672 | P1672_mbaA_E553A_homol2_R | ccgggagagctcgatatcgcatgcggtacctctaggatggtggcgggttaaaagg |
| P1553 | P1553_itac_mbaa_pdest17_f | tgtcgtactaccatcaccatcaccatcacctcgaagcggccgcagagaacctg-tattttcagggtatggtgatcaatccgatcctgcatt |
| P1554 | P1554_itac_mbaa_pdest17_r | gcttcctttcgggctttgttagcagcctcgaatcactcgagaggcacttctaacggcattc |
| P1620 | P1620_MBP_ITAC_F | gaaataattttgtttaactttaagaaggagatatacatatgggcagcagccatcatcatc |
| P1621 | P1621_MBP_ITAC_R | ggattgatcaccataccctgaaaatacaggttctcgcggccgcctgcgcgtctttcagggct |
| P1648 | P1648_pOML213_mbaA_E576K_R | caatccagtttgctttgcaattgg |
| P1649 | P1649_pOML213_mbaA_E576K_F | tttggcacgatcgaccgctg |
| P1650 | P1650_pOML213_seq_F | agactgtcgatgaagccctgaaa |
| P1660 | P1660_pOML213_mbaA_L618V_F | tcagcgaattgacttctgccg |
| P1661 | P1661_pOML213_mbaA_L618V_R | aattggctcaatttatccatcgacaggc |
| P1662 | P1662_pOML213_mbaA_E483K_F | cagcaatttcttaatgtgttcgcc |
| P1663 | P1663_pOML213_mbaA_E483K_R | ctcaatgccgataccgccatgt |
| P1667 | P1667_pOML213_mbaA_E553A_F | gcagcaatactgcgaagccc |
| P1668 | P1668_pOML213_mbaA_E553A_R | gctggcaatctgaattgctcgg |

**Table S3. Biofilm formation of wild-type and evolved populations of *V. cholerae***

| **Populations** | **Biofilm formation** $OD_{595}$[a] | **Relative biofilm formation** | ***P* values**[b] | ***N***[c] |
|---|---|---|---|---|
| Parental *V. cholerae* | 0.68 ± 0.09 | 1 | - | 3 |
| Planktonic-evolved population1 | 1.11 ± 0.14 | 1.69 ± 0.30 | 0.9991 | 3 |
| Planktonic-evolved population 2 | 1.80 ± 0.55 | 2.85 ± 1.03 | 0.8227 | 3 |
| Biofilm-evolved population 1 | 6.32 ± 1.12 | 9.73 ± 2.30 | 0.0002 | 3 |
| Biofilm-evolved population 2 | 6.68 ± 0.92 | 10.21 ± 1.96 | <0.0001 | 3 |
| Biofilm-evolved population 3 | 6.40 ± 0.20 | 9.83 ± 1.34 | 0.0002 | 3 |
| Biofilm-evolved population 4 | 7.52 ± 1.02 | 11.37 ± 1.89 | <0.0001 | 3 |
| Biofilm-evolved population 5 | 5.16 ± 0.98 | 8.04 ± 2.16 | 0.0023 | 3 |
| Biofilm-evolved population 6 | 5.18 ± 0.44 | 7.82 ± 0.86 | 0.0022 | 3 |

[a] Absorbance measured at 595 nm for dissolved biofilms stained with crystal violet (CV) in 96-well plates.
[b] One way ANOVA to test for differences in biomass formation compared to wild-type (α=0.05) and followed by Dunnett test to correct for multiple testing; F (DFn, DFd) 0.6042, (8, 18), overall *P* value <0.0001.
[c] Sample size (biological replicates) tested to determine biofilm formation
Errors are reported as the standard error of the mean

**Table S4. P-value table from one-way ANOVA test comparing biofilm formation of all mbaA variants**

| | Wild-type | G425A | ΔmbaA | ΔEAL | E553A | G425A /E576K | E576K | E483K | L618V |
|---|---|---|---|---|---|---|---|---|---|
| Wild-type | n.a. | n.a. | n.a. | n.a. | n.a. | n.a. | n.a. | n.a. | n.a. |
| G425A | 0.1459 | n.a. | n.a. | n.a. | n.a. | n.a. | n.a. | n.a. | n.a. |
| ΔmbaA | <0.0001 | <0.0001 | n.a. | n.a. | n.a. | n.a. | n.a. | n.a. | n.a. |
| ΔEAL | 0.0085 | 0.0317 | 0.8950 | n.a. | n.a. | n.a. | n.a. | n.a. | n.a. |
| E553A | 0.0003 | 0.0012 | 0.7435 | >0.9999 | n.a. | n.a. | n.a. | n.a. | n.a. |
| G425A/E576K | 0.0002 | 0.0004 | >0.9999 | 0.9995 | 0.9971 | n.a. | n.a. | n.a. | n.a. |
| E576K | <0.0001 | <0.0001 | <0.0001 | <0.0001 | <0.0001 | <0.0001 | n.a. | n.a. | n.a. |
| E483K | <0.0001 | <0.0001 | 0.0407 | 0.0020 | 0.0007 | 0.0083 | 0.6400 | n.a. | n.a. |
| L618V | <0.0001 | <0.0001 | 0.0232 | 0.0013 | 0.0004 | 0.0055 | 0.7721 | >0.9999 | n.a. |

#### 2. Supplementary figures

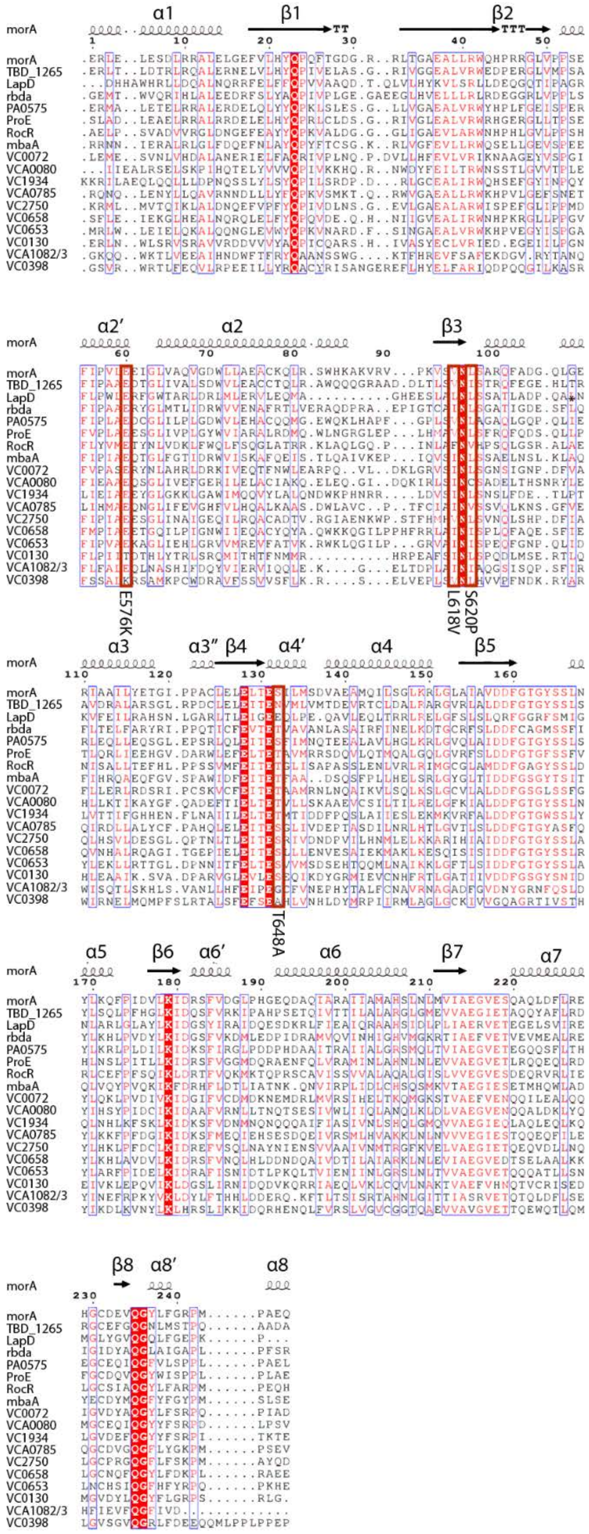


**Figure S1: Alignment of selected EAL-domains from GGDEF-EAL proteins**

Multiple sequence alignment of putative conserved EAL-domains from *V. cholerae* C6706 as well as EAL-domains with solved crystal structures. The secondary structure based on the crystal structure of morA (PDB ID: 4RNH) is depicted on the top of the figure [6]. α-turns and β-turns are depicted as TTT and TT, respectively. Mutations selected for by biofilm evolution in mbaA are annotated and marked by a red box. Conserved residues defined as residues with a similarity global score >0.7 are depicted as red characters on a white background with a blue box around. Strictly conserved residues are rendered as white characters on a red background with a blue box.

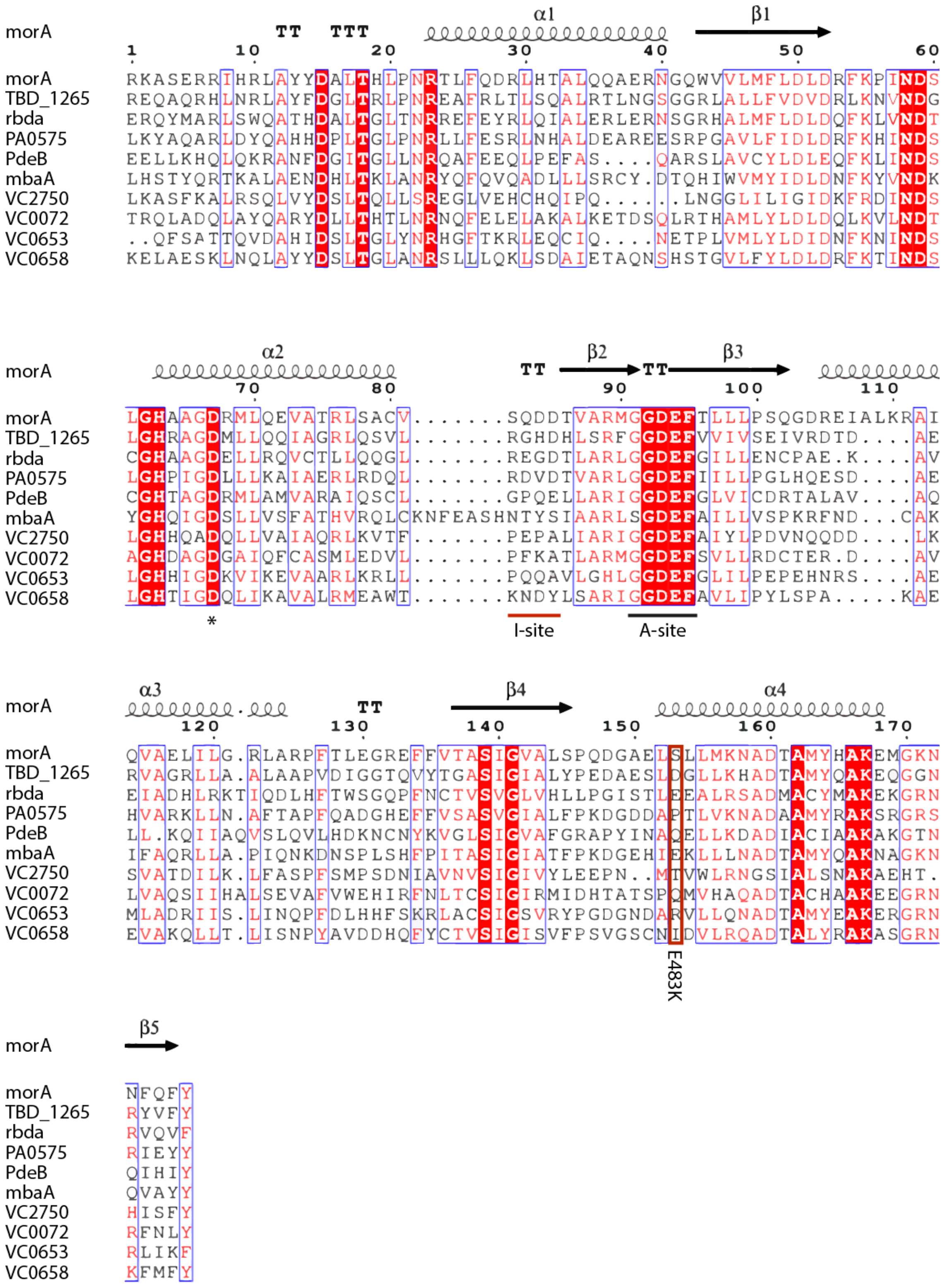


**Figure S2: Alignment of selected GGDEF-domains from GGDEF-EAL proteins**

Multiple sequence alignment of putative conserved GGDEF-domains from *V. cholerae* C6706 as well as GGDEF-domains with solved crystal structures. The secondary structure based on the crystal structure of morA (PDB ID: 4RNH) is depicted on the top of the figure [6]. A-turns and β-turns are depicted as TTT and TT, respectively. Mutations selected for by biofilm evolution in mbaA are annotated and marked by a red box. Conserved residues defined as residues with a similarity global score >0.7 are depicted as red characters on a white background with a blue box around. Strictly conserved residues are rendered as white characters on a red background with a blue box.

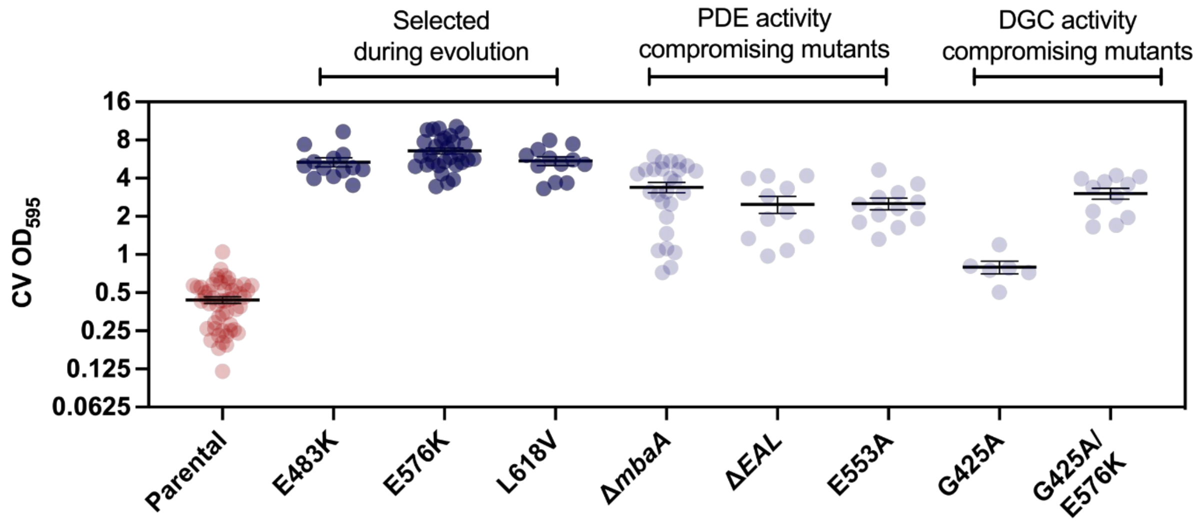


**Figure S3: Biofilm formation of parental *V. cholerae* and all tested MbaA mutants**

Biofilm formation of parental *V. cholerae* and all tested MbaA mutants. Biofilm production was quantified by measuring the optical density at 595 nm of dissolved biofilms stained with crystal violet (CV). Each dot represents one biological replicate. Error bars represent standard error of the mean

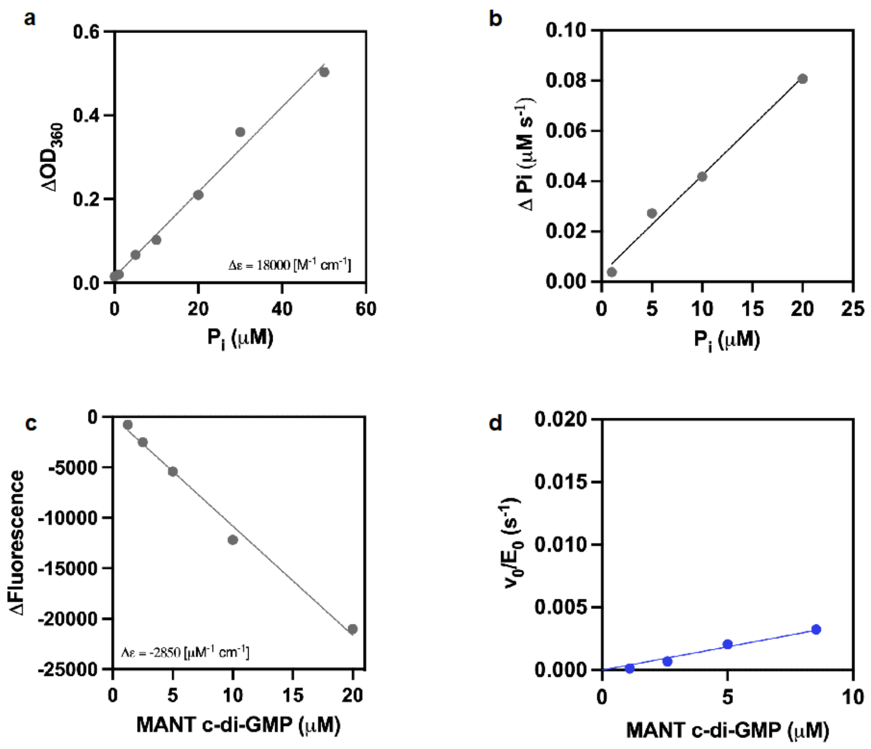


**Figure S4: Validation of measurements of DGC and PDE activity**

**a.** Determination of the extinction coefficient for the reaction of $P_i$ with MESG mediated by purine nucleoside phosphorylase (PNP) that was used to measure DGC activity. The reaction was measured as changes in $OD_{360}$ before and after incubation with purine nucleoside phosphorylase (PNP). **b.** Reaction velocity of pure $P_i$ with MESG with substrate concentrations ranging from 1.25 to 20 µM. **c.** Determination of extinction coefficient for the hydrolysis of MANT-c-di-GMP using emitted fluorescence of MANT-c-di-GMP before and after incubation with wild-type MbaA **d.** MANT-c-di-GMP hydrolysis curves for E553A with substrate concentrations ranging from 1.25 to 8 µM. E553A displayed non-saturation kinetics and a 5.3-fold decrease in $k_{cat}/K_M$, indicating that impaired substrate affinity contributed to the decrease in $k_{cat}/K_M$ compared to wild-type MbaA (Tab. 3).
